# The putative Type 4 secretion system effector BspD is involved in maintaining envelope integrity of the pathogen *Brucella*

**DOI:** 10.1101/2024.03.18.585630

**Authors:** Maren Ketterer, Petra Chiquet, Jaroslaw Sedzicki, Maxime Québatte, Christoph Dehio

## Abstract

Brucellosis is a debilitating disease caused by the Gram-negative, facultative intracellular zoonotic pathogen *Brucella*. *En route* to its intracellular replicative niche, *Brucella* encounters various stressful environments that may compromise envelop integrity. Here we show, that the proposed Type 4 secretion system (T4SS) effector BspD is a conserved protein of the Rhizobiales, which does not show signs of co-evolution with the presence of a T4SS or a certain lifestyle. We further present data indicating that BspD is critical for *Brucella abortus* envelope integrity in the stationary phase and in the presence of EDTA, a compound known to destabilize the outer membrane. Deletion of *bspD* resulted in abnormal bacterial morphologies, indicating its involvement in maintaining envelope integrity. Additionally, the absence of BspD led to the formation of fewer and smaller intracellular microcolonies in a macrophage infection model. From our observations, we propose that BspD of *B. abortus* is critical for preserving the integrity of the bacterial envelope, particularly under stressful conditions, which may enhance *Brucella*’s ability to survive within host cells.

**Importance:** Brucellosis, caused by the intracellular pathogen *Brucella*, poses a significant health threat. Understanding how *Brucella* adapts to stressful environments is crucial. This study unveils BspD, a conserved protein within the Rhizobiales order, as a key player in maintaining *Brucella*’s envelope integrity. Remarkably, BspD’s function appears independent of the presence of a Type 4 Secretion System (T4SS) or specific lifestyle. Deletion of *bspD* resulted in compromised envelope integrity, abnormal bacterial morphologies, and reduced intracellular microcolony formation. These findings underscore BspD’s critical role, particularly in stressful conditions like the stationary phase and EDTA exposure, and highlight its significance in *Brucella*’s survival within host cells. This elucidation deepens our understanding of *Brucella* pathogenesis and may inform future therapeutic strategies against brucellosis.

## Introduction

The Rhizobiales, an order of diverse Gram-negative α-Proteobacteria, are specialized to a wide range of niches; spanning from free living bacteria to plant symbionts and intracellular mammalian pathogens (1, 2). The genus *Brucella* comprises the facultative intracellular zoonotic pathogens *Brucella abortus*, *Brucella suis* and *Brucella melitensis*, which are the causative agents of Brucellosis, a severe zoonosis manifesting through a broad range of symptoms, encompassing undulant fever and flu-like episodes (3).

As a facultative intracellular pathogen *Brucella* has evolved strategies to survive and replicate within host cells. Following uptake, the *Brucella*-containing vacuole (BCV) is trafficking along the canonical endocytic route to at least the late endosomal stage (4–8). Acidification of the BCV serves as signal for triggering expression of the bacterial Type 4 Secretion System (T4SS) VirB, which then facilitates rerouting of the BCV to the endoplasmic reticulum (ER) where bacteria establish their replicative niche (4, 9, 10, 7, 8, 11). T4SSs are pivotal virulence factors of numerous host-interacting bacteria; they represent large macromolecular complexes spanning the bacterial envelope that facilitate the export of cargo from the bacterial cytoplasm, such as the translocation of toxins to the extracellular environment or of bacterial effector proteins directly into host cells (reviewed in (12)). Deletion of any gene encoding an essential component of the VirB machinery incapacitates the mutant bacteria to reach the ER and to replicate intracellularly, demonstrating the essential role of the T4SS in *Brucella*’s intracellular lifestyle (7, 13). In contrast, none of the growing list of putative T4SS effector proteins has been found to be individually essential for intracellular replication or survival (14). Despite the ample evidence for a central role of the VirB T4SS for *Brucella*’s virulence (15–17), we are still lacking a molecular understanding of how the machinery works, including the nature of translocation motif(s) and recruitment mechanism(s) of the effectors and the process of their translocation.

Before reaching the ER-derived replication niche, bacteria trafficking within the BCV encounter a challenging and ever changing environment, including acidification, the action of cationic peptides and lytic enzymes, and the activity of reactive oxygen species (18, 19). A key component to withstand these chemical stressors is *Brucella*’s cell envelope, which consists of an inner membrane, an outer membrane, and the periplasmic space with the peptidoglycan layer in between. *Brucella* is long known for its remarkable resistance to chelators, detergents, and cationic peptides (20, 21), however, we are only now beginning to gain insights into the molecular mechanisms facilitating stress resistance and maintenance of envelope integrity. Some of the involved components are the periplasmic proteins EipA (22) and EipB (23), the TamB homologue MapB (24), and *Brucella*’s distinct LPS (21, 25). They were shown to confer resistance to several cell wall stressors, such as hyperosmotic and acidic environments, cell wall-active antibiotics, metal chelators, and cationic peptides, and are important for survival within the host (21–25).

Interestingly, the putative T4SS effector protein BspD (BAB1_1611) of unknown function (26, 27) is encoded on chromosome 1 directly downstream from the gene encoding the envelope integrity protein EipA (22). This locus architecture prompted us to test for a potential role of BspD in envelope integrity. Through a combination of *in silico* analysis, *in vitro*, and *in cellulo* experiments, we show that (i) BspD is a conserved protein confined to the Rhizobiales; (ii) the deletion of *bspD* confers a mild, but significant growth defect in rich broth in stationary phase and (iii) the *bspD* mutant displays sensitivity to EDTA. We conclude that BspD has a conserved function in envelope homeostasis that is likely independent of any T4SS-related function. Further, we show that deletion of *bspD* compromises intracellular microcolony formation in macrophages, highlighting the role of BspD in mediating stress tolerance *in cellulo*, but this mutant phenotype could also reflect the proposed role of BspD as a T4SS effector.

## Results

### The putative T4SS effector BspD is conserved in the *Rhizobiales*

In *Brucella*, the gene encoding the putative T4SS effector BspD is located directly downstream to the genes encoding the envelope integrity protein EipA and the essential cell cycle response regulator CtrA and its upstream regulator, the histidine phosphotransferase ChpT (22), as well as PicC, an essential protein of unknown function (14) (Fig. 1A). Several genes surrounding *bspD* have CtrA-binding sites (28), and are essential for growth of *Brucella* on rich medium (14) (Fig. 1A). Prompted by the conserved synteny (Fig. 1A) we wanted to explore the evolutionary history of BspD in more detail. Therefore, we analysed the presence of the *bspD* gene and conservation of the microsynteny with *eipA*, *chpT* and *ctrA* within the entire Alphaproteobacteria. Using the OMA browser (29, 30) to search for *bspD* orthologues, *bspD* was found to be confined to the Rhizobiales and conserved in a majority of its member genera and species (Fig. 1B). Moreover, its presence does not correlate with the presence of a VirB T4SS or a particular lifestyle. For instance, BspD homologues are found in many soil-dwelling species such as *Methylocella silvestris* and *Strakeya novella*, as well as classical plant symbionts such as *Rhizobium meliloti* and *Mesorhizobium japonicum*, while the mammalian pathogens comprised by the *Bartonellae* do not encode BspD (Fig. 1B). Other than *bspD*, the genes coding for EipA, CtrA, and ChpT are not confined to the Rhizobiales and are also found in the Sphingomonadales, Caulobacterales, Parvulaculales, and the Rhodobacterales (22). Within species of the Rhizobiales, the microsynteny of *bspD* with at least *eipA, chpT*, and *ctrA* extends beyond the genus *Brucella* also to species of the *Phyllobacteriaceae* and *Rhizobiacea*, such as *Mesorhizobium japonicum* and *Rhizobium meliloti*, respectively (Fig. 1A, B), while in other members of the Rhizobiales the microsynteny is limited to *eipA* (Fig. 1B) (31). A protein sequence alignment further revealed that BspD of *Brucella* and *Ochrobactrum* is truncated compared to orthologs of other species (Fig. S1), which coincides with the *Brucellaceae* forming a robust cluster in a protein-based phylogenetic analysis (Fig. S2).

**Figure 1:**
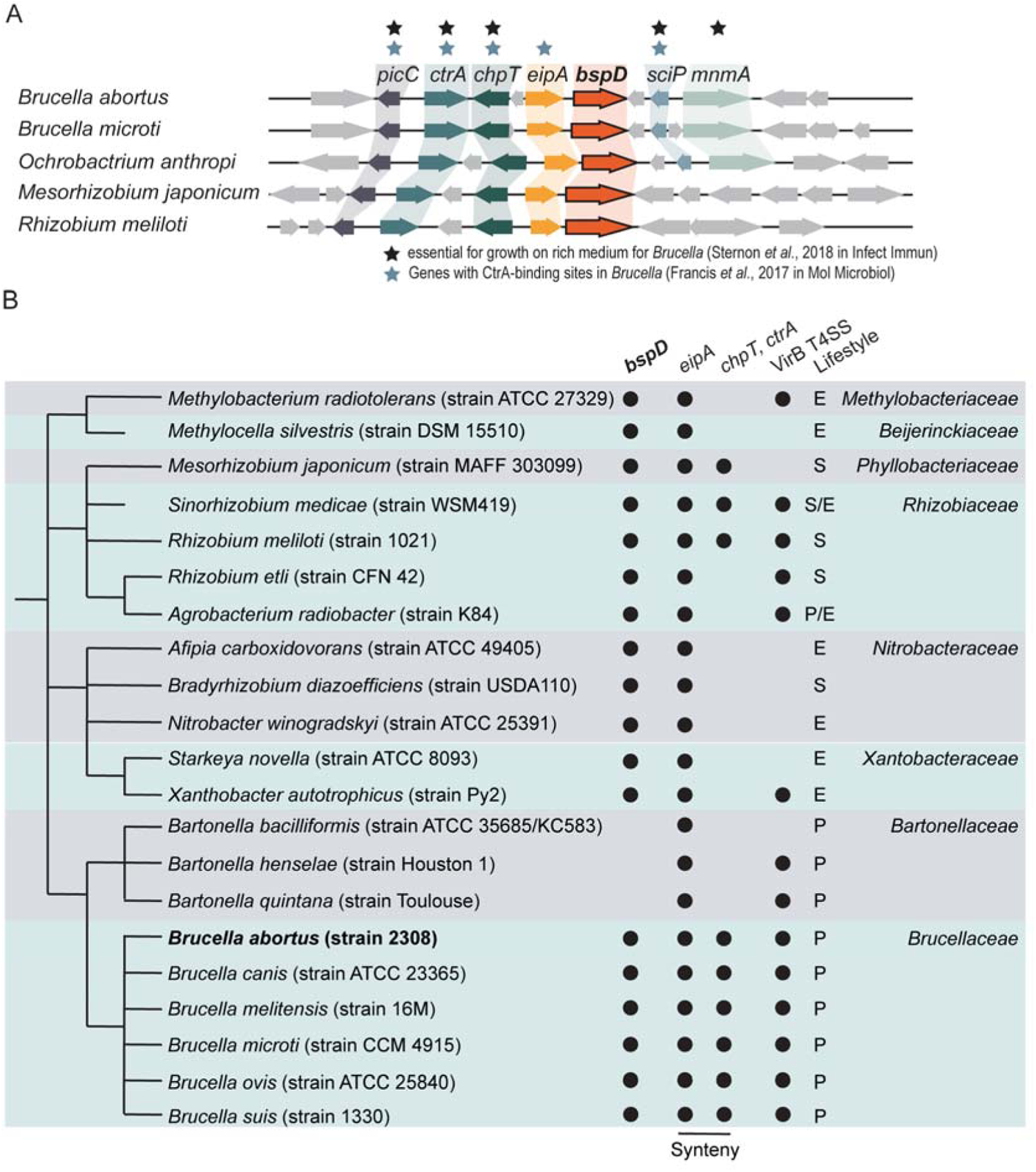
The putative effector BspD is conserved in the Rhizobiales. **(A)** *bspD* is co-conserved with *picC*, *ctrA*, *chpT*, and *eipA* in a subset of the Rhizobiales. Gene synteny redrawn from SyntTax. Black stars depict genes reported to be essential for growth on rich medium for *Brucella*. Blue stars mark genes with reported CtrA-binding sites in *Brucella*. **(B)** Simplified phylogenetic tree of the genus Rhizobiales based on the species phylogeny of hierarchical orthologous groups (HOGs) as defined by the OMA browser. Presence of *bspD* (BAB1_1611) and *eipA* (BAB1_1612) orthologs based on their respective HOGs is indicated with a filled circle. Co-conservation with *ctrA* and *chpT* in the same locus is depicted with a filled circle deduced from SyntTax. Presence of VirB T4SSs is depicted with a filled circle as inferred from the SecReT4 database. The different lifestyles (S – Plant associated, P – Animal Pathogen, E – Environmental) were inferred from literature.

### Deletion of *bspD* leads to growth defect in rich broth

Given the conserved linkage of *bspD* with genes encoding the envelope integrity protein EipA (22) or the essential growth regulators CtrA/ChpT (14, 28), we aimed at testing whether BspD might have a conserved function in growth control or envelope homeostasis.

To accomplish this, we generated a Δ*bspD* deletion mutant, and employed a previously described miniTn7-based strategy for integration of a single chromosomal copy of a *bspD* rescue construct between *glmS* and *recG* of chromosome 1 (26) (Fig. 2A). Considering the possibility that *eipA* and *bspD* might constitute a bi-cistronic operon co-transcribed from the upstream located CtrA-controlled *eipA* promoter (22), we integrated a fragment encoding both genes, including the *eipA* promoter (Δ*bspD::eipA-bspD*), to ensure native expression of BspD. To eliminate the potential influence of an additional copy of *eipA*, we also constructed an isogenic Δ*bspD* mutant strain containing an insertion solely for *eipA* along with its promoter region (Δ*bspD::eipA*) (Fig. 2A).

**Figure 2:**
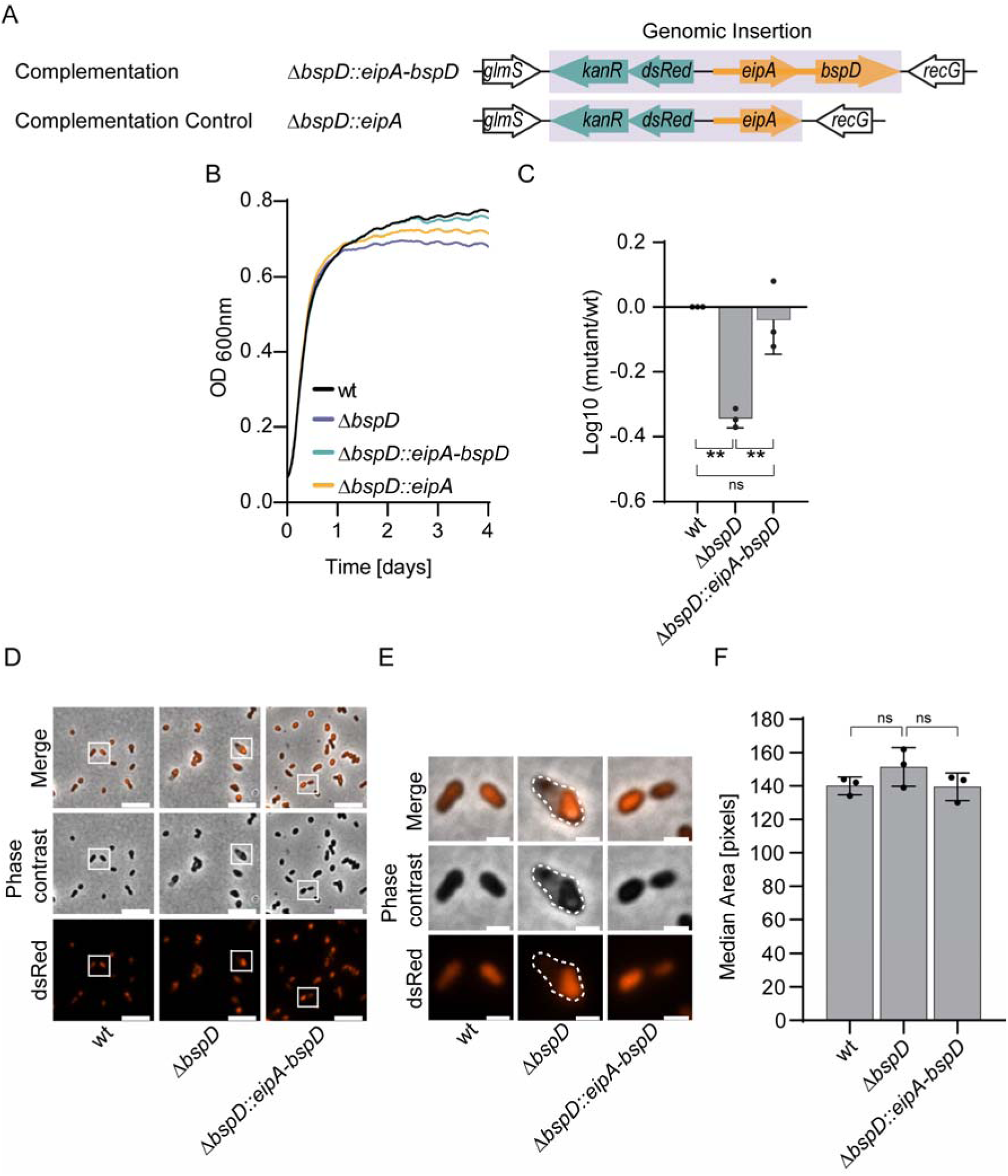
Deletion of *bspD* results in a stationary phase survival defect of *Brucella*. **(A)** Overview of Δ*bspD* complementation strategy. To ensure native expression levels the 5’ non-coding regions of *eipA* and *bspD* (orange thick line) encoding potential promoters were included. Violet shaded area – genomic insertion. *kanR* – Kanamycin resistance cassette under *P_aphT_*. *dsRed* – *dsRed* under *P_aphT_*. Black outline – genomic context. Teal – *kanR* and *dsRed* genes. Orange – *eipA* and *bspD* with 5’ non-coding regions. **(B)** Optical density of indicated *Brucella* strains grown in 150 µl TSB in a 96 well plate over 96 h with a starting OD_600nm_ of 0.08. The OD_600nm_ was measured every 10 min in a Synergy H1 plate reader. Lines depict means of 3 independent experiments with a 2^nd^ order smoothing of 15 neighbors applied in GraphPad Prism 9.3.1. **(C)** Comparison of recovered CFU of wild-type, deletion mutant and complemented mutant from (B) at 96 h. Cultures were serially diluted and plated on TSA plates. After 3-4 days of incubation at 37°C CFUs were enumerated and the Log10 survival was calculated by deletion of mutant CFU ml^−1^ by wt CFU ml^−1^. Corresponding CFU enumerations are displayed in Fig. S3A. **(D)** Phase contrast and fluorescence micrographs of wt, deletion mutant, and rescue cultured in TSB for 96 h before fixation. Insets in Δ*bspD* highlight bacterial cells with plasmolysis. Scale bar = 5 μm. Representative images of 3 independent replicates. **(E)** Enlarged insets of (D) showing bacteria with apparent plasmolysis. Broken line = outline of bacterium. Scale bar = 1 μm. **(F)** Evaluation of bacterial morphology of bacteria recovered from (B) at 96 h. The area of the dsRed signal of >500 single bacteria from 3 independent experiments was measured using CellProfiler as a proxy for bacterial size. Displayed are the median areas with standard deviations.

We then tested the growth of the various isogenic *B. abortus* strains in TSB over 96 h in comparison to the parental wild-type strain. Compared to wild-type and the rescued mutant strain Δ*bspD::eipA-bspD*, the Δ*bspD* mutant and Δ*bspD::eipA* control strain displayed a growth defect in rich broth characterized by an earlier entry into stationary phase (Fig. 2B). CFU counts performed at the end of the experiment showed that mutant levels were approximately 50% lower than wild-type counts (Fig. 2C, Fig. S3A). Notably, a Δ*virB9* mutant displayed comparable growth to wild-type bacteria (Fig. S4), suggesting that the observed growth defect of the Δ*bspD* mutant is independent of a functional T4SS.

Although we did not observe significant morphological changes, such as alterations in cell size (Fig. 2D-F), we did observe heightened levels of plasmolysis in the Δ*bspD* mutant. This was indicated by the contraction of the dsRed-labelled cytoplasm, as depicted in Fig. 2D and E.

### The **Δ***bspD* mutant is destabilized by EDTA

Next we investigated if the Δ*bspD* mutant might be compromised in envelope homeostasis using the same conditions as previously described for the Δ*eipA* mutant (22). To this end we tested the sensitivity of the Δ*bspD* mutant vs. wild-type to EDTA, high NaCl concentration, and ampicillin. The Δ*bspD* mutant displayed significant sensitivity to EDTA in the stress plate assay when compared to growth in absence of any stressor, but only mild sensitivity to high NaCl concentrations (Fig. 3A, Fig. S3B). There was no difference in the sensitivity of the Δ*bspD* mutant to ampicillin (Fig. 3A, Fig. S3B).

**Figure 3:**
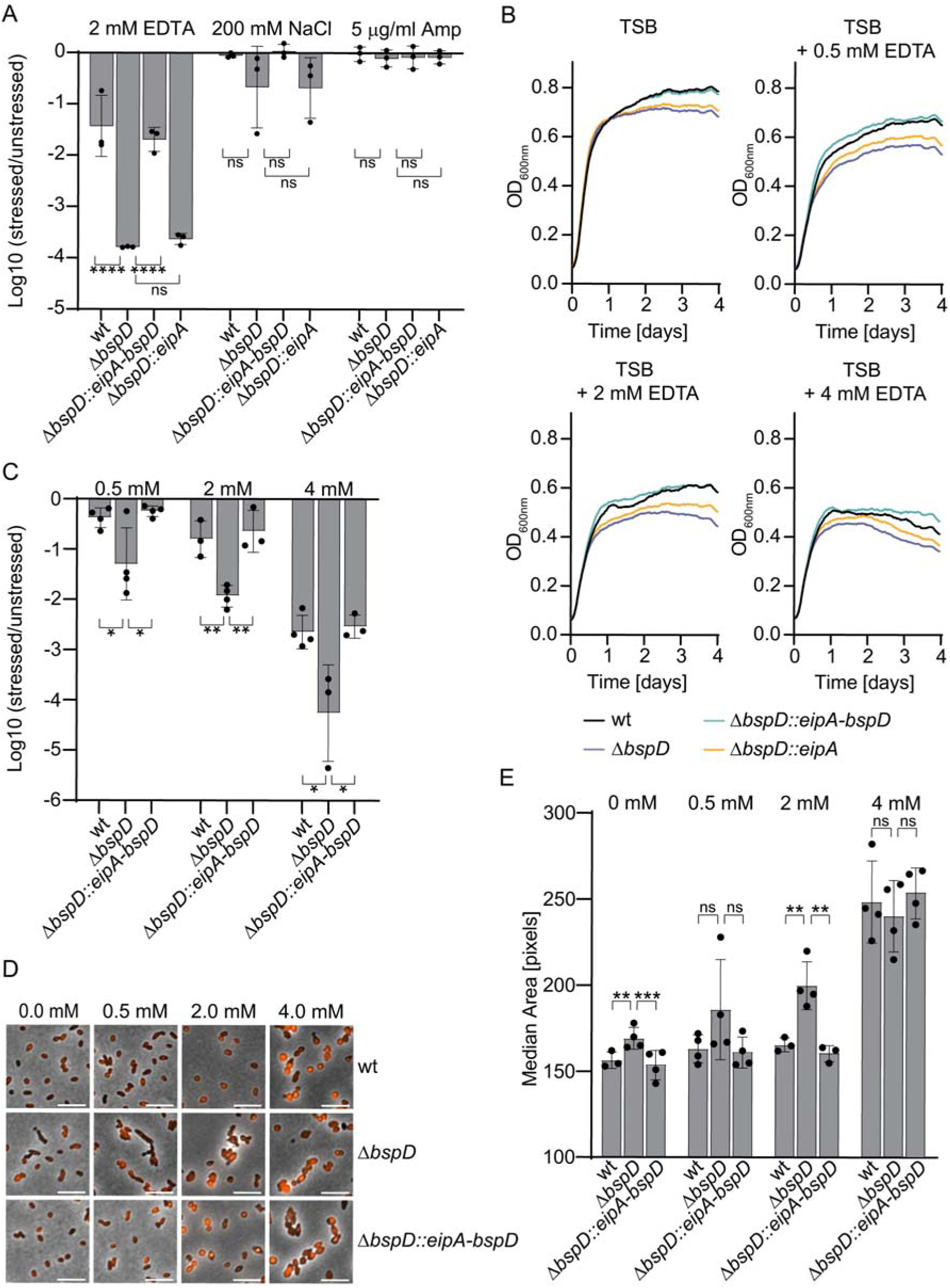
Deletion of *bspD* leads to a survival defect in presence of EDTA. (A) Plate stress assay to assess survival in presence of envelope stressors. Indicated *Brucella* strains were grown to stationary phase and the density was adjusted to 1×10^8^ CFU ml^-1^. Samples were serially diluted, and spotted onto plain TSA plates or TSA plates supplemented with either 2 mM EDTA, 200 mM NaCl, or 5 µg ml^−1^ ampicillin. CFUs were enumerated after at least 3 days of growth at 37°C. Y-axis shows for each strain the log10 reduction for the ratio of CFUs on supplemented TSA versus CFUs on plain TSA. Dots represent means of 2 technical replicates of 3 independent experiments ±SD. Corresponding CFU enumerations are displayed in Fig. S3B. **(B)** Optical density of indicated *Brucella* strains grown in 150 µl plain TSB or TSB supplemented with either 0.5, 2, or 4 mM EDTA in a 96 well plate over 96 h with a starting OD_600nm_ of 0.08. The OD_600nm_ was measured every 10 min in a Synergy H1 plate reader. Depicted is one representative experiment of 4 independent replicates. Data is displayed with a 2^nd^ order smoothing of 15 neighbors applied in GraphPad Prism 9.3.1. **(C)** Relative survival of mutants vs wt bacteria recovered at 96 h from (B). Samples were serially diluted and plated on TSA plates. After 3-4 days of incubation at 37°C CFUs were enumerated and the Log10 survival was calculated by deletion of mutant CFU ml^−1^ by wt CFU ml^−1^. Outliers were removed as described in the methods section. Corresponding CFU enumerations are displayed in Fig. S3C. **(D)** Representative micrographs of bacteria recovered from (B). Red – dsRed, black – phase contrast. Scale bar = 5 μm. **(E)** Evaluation of bacterial morphology of bacteria recovered from (B) at 96 h. The area of the dsRed signal of >500 single bacteria from 4 independent experiments was measured using CellProfiler as a proxy for bacterial size. Displayed are the median areas measured in the independent experiments ±SD. Outliers were removed as described in the methods section.

To further characterize the nature of the increased sensitivity of the Δ*bspD* mutant to EDTA we grew bacteria in rich broth in the presence of increasing EDTA concentrations for 4 days continuously measuring the OD_600nm_, and evaluating the CFU ml^−1^ and morphology at endpoint (Fig. 3B-E). We observed that the stationary phase defect of the Δ*bspD* mutant in TSB was exacerbated in presence of low EDTA concentrations (Fig. 3B), which correlated with a survival defect based on CFU counting at the endpoint (Fig. 3C, Fig. S3C). We further observed that the bacterial shape and size of wild-type bacteria was stable from 0-2 mM EDTA, while the size of the Δ*bspD* mutant increased steadily and showed aberrant morphologies with increasing EDTA concentrations (Fig. 3D and E), indicating a destabilization of the bacterial envelope. These findings suggest that BspD is required for envelope integrity in stationary phase and under outer membrane stress conditions. Notably, the outcomes we observed in the presence of hyperosmotic stress or ampicillin do not indicate a broad destabilization in the Δ*bspD* mutant. Instead, they imply a more specific role of BspD in envelope integrity.

### The **Δ***bspD* mutant is compromised in the formation and growth of intracellular microcolonies

Given the observed extracellular stationary phase and outer membrane stress defect of the Δ*bspD* mutant, we thought to determine if this mutant’s ability to reach the intracellular replication niche and then form intracellular microcolonies was compromised. To achieve this, we infected RAW264.7 macrophages and evaluated the proportion of cells containing microcolonies (Fig. 4A and B), as well as the size of these microcolonies after 27 hpi using CellProfiler for automated image analysis (Fig. 4A and C). To this end we established a CellProfiler pipeline as previously described (32) and detailed in the method section. In brief, thresholds for identifying microcolonies were determined by comparing the size of infection sites of the replication-deficient Δ*virB9* strain with microcolonies formed by wild-type *B. abortus* in RAW264.7 macrophages after 27h of infection (Fig. S5). Using this pipeline, we observed a significant decrease in the proportion of host cells with microcolonies upon deletion of *bspD* (Fig. 4A and B). Furthermore, the microcolonies formed were smaller compared to wild-type microcolonies (Fig. 4A and C). These findings support our hypothesis that BspD may play a crucial role in envelope homeostasis, thereby ensuring survival in adverse environments. However, we cannot exclude the possibility that a putative effector function of BspD also contributes to the observed intracellular phenotype.

**Figure 4:**
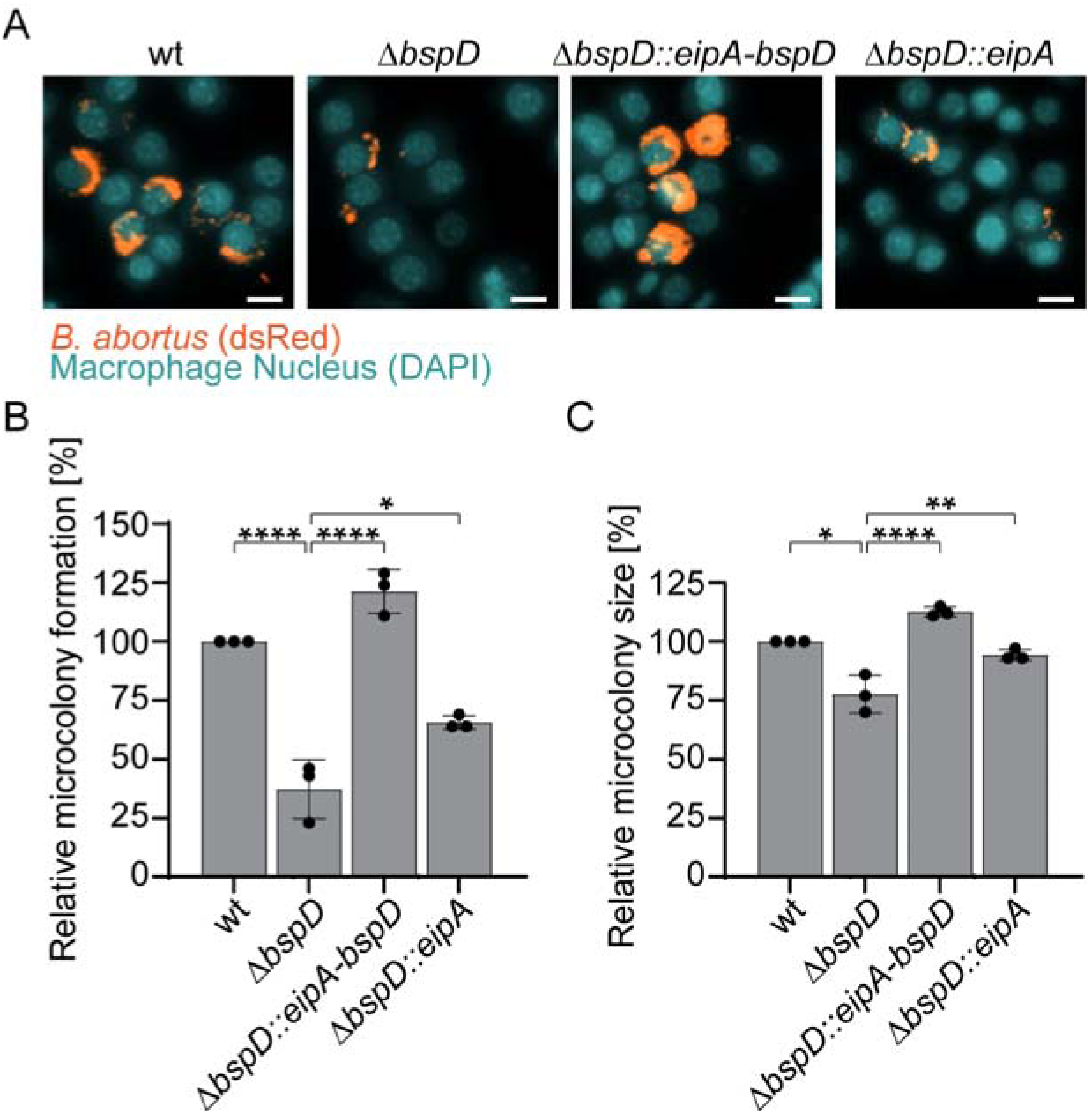
The Δ*bspD* mutant is compromised in intracellular replication within macrophages. RAW264.7 macrophages were infected with an MOI of 50 for 27 h. After fixation and staining of host cell nuclei with DAPI, samples were imaged using a Molecular Devices ImageXpress microscope. n=3. **(A)** Representative images of infected RAW264.7 macrophage-like cells. Cyan – host cell nuclei. Orange – dsRed-expressing *Brucella*. Scale bar = 10 μm. **(B)** Relative microcolony formation of different *Brucella* strains normalized to wt. Relative microcolony formation corresponds to the quantity of host cells with identified microcolonies. Individual replicates are shown as dots. **(C)** Relative microcolony size formed by different *Brucella* strains expressed as percentage relative to wild-type level. The relative microcolony size of the mutants based on the median size of about 7000 identified infection sites in each experiment was calculated in relation to the wt. Dots represent individual experiments ±SD.

## Discussion

To establish infection as an intracellular pathogen, *Brucella* must overcome two significant challenges: firstly, it must endure various multi-stress environments, and secondly, it must manipulate the host cell to ensure its survival and replication. In this study, we present data suggesting a role for the putative T4SS effector BspD in envelope homeostasis in *Brucella*, contributing to stress resistance and survival in axenic conditions and within host cells.

BspD is preserved across the Rhizobiales species with varying lifestyles, including those with and without T4SSs, except for pathogens in the *Bartonella* genus, which lack BspD, likely due to gene loss in the process of massive genome reduction in their evolutionary trajectory to blood-borne pathogens (33). This suggests that the *bspD* genes in the Rhizobiales are true orthologues derived from a common ancestor, indicating an ancient function of BspD that doesn’t depend on the T4SS. Indeed, our observations indicate that the deletion of *bspD* results in a mild stationary phase defect. Stationary phase conditions can entail stresses such as acidification and the accumulation of reactive oxygen intermediates (18, 34). Our observations show further that the Δ*bspD* mutant experiences a notable reduction in fitness when exposed to EDTA on agar. EDTA disrupts the outer membrane by removing divalent cations that help stabilize it via interactions with the hydrophilic part of LPS (20, 35), suggesting BspD plays a role in enhancing the outer membrane’s resistance to stress. However, this mutant doesn’t show significant fitness loss under osmotic stress with NaCl or peptidoglycan stress with ampicillin, unlike the *eipA* gene deletion, which affects fitness under all three stress conditions (22). This indicates that while *eipA* and *bspD* may be part of the same operon, their proteins serve different, only somewhat overlapping roles in managing envelope stress.

The bspD deletion mutant exhibited morphological changes when exposed to EDTA, related to those previously observed in *Escherichia coli*, including increased size and plasmolysis (36). Several genes near *bspD* contain CtrA-binding sites, such as *picC*, *ctrA*, *chpT*, *sciP*, and *mnmA* (28). These genes are crucial for *Brucella*’s growth on rich media (14) and are linked to cell cycle regulation and outer membrane composition (37, 28, 38). This suggests BspD may influence LPS synthesis or modification, potentially explaining *Brucella*’s notable EDTA resistance (20, 21).

Besides EDTA sensitivity, the *bspD* deletion mutant showed intracellular impairments, forming fewer and smaller microcolonies than wild-type *Brucella*. This underscores BspD’s role in tolerating intracellular stress. Given BspD’s possible function as a T4SS effector, it’s important to consider that its deletion might delay intracellular trafficking, leading to the observed smaller microcolonies. Identifying the precise reasons for these intracellular anomalies should be a priority for future studies.

## Material and Methods

### Bacterial strains and growth conditions

All manipulations involving live *Brucella abortus* 2308 and its derivatives were conducted in the biosafety level 3 facility at either the Swiss Tropical and Public Health Institute or the Biozentrum of the University of Basel, both located in Basel, Switzerland. These manipulations were carried out following standard operational procedures (SOPs).

Bacterial strains used in this study are listed in Table 1. *E. coli* strains were cultivated in lysogeny broth (LB) or on agar plates (LA) at 37°C overnight supplemented with appropriate antibiotics.

**Table 1:** Strains used in this study.

| Strain | Relevant characteristics | Reference/Source | Identifier/Description |
| --- | --- | --- | --- |
| ***Escherichia coli*** |  |  |  |
| Stellar | *F*<sup>-</sup>, *endA1*, *supE44*, *thi-1*, *recA1*, *relA1*, *gyrA96*, *phoA*, $\Phi 80d$ *lacZ*Δ M15, $\Delta$(*lacZYA-argF*) U169, $\Delta$(*mrr-hsdRMS-mcrBC*), Δ*mcrA*, λ- | Takara bio | Standard cloning strain |
| JKE201 | MFDpir Δ*mcrA* $\Delta$(*mrr-hsdRMS-mcrBC*) *aac(3)IV::lacIq* | (45) | Donor strain for conjugation |
| ST18 | *pro*, *thi*, *hsdR*<sup>+</sup>, T<sub>p</sub><sup>r</sup>, Sm<sup>r</sup>; chromosome::*RP4-2 Tc::Mu-Kan::Tn7/*λ*pir*, Δ*hemA* | (46) | Donor strain for conjugation of miniTn7 |
| JSE273 | JKE201 pTNS2 | This study | Helper strain for insertion of miniTn7 |
| ***Brucella abortus*** |  |  |  |
| RCBr01 | *B. abortus* 2308 | J. P. Gorvel, Centre d'Immunologie de Marseille-Luminy (CIML) |  |
| RCBr03 | *B. abortus* 2308 Δ*virB9* | (47) | |
| MQBr19 | *B. abortus* 2308 Δ*bspD* | This study | |
| MKBr08 | *B. abortus* 2308 (*tn7*)::*P*<sub>tet</sub>:empty | This study |  |
| MKBr09 | *B. abortus* 2308 Δ*bspD* (*tn7*)::*P*<sub>tet</sub>:empty | This study | |
| MKBr26 | *B. abortus* 2308 Δ*virB9* (*tn7*)::*P*<sub>tet</sub>:empty | This study | |
| MKBr34 | *B. abortus* 2308 Δ*bspD* (*tn7*)::*eipAbspD* | This study | |
| MKBr40 | *B. abortus* 2308 Δ*bspD* (*tn7*)::*eipA* | This study | |

*Brucella* strains were grown in tryptic soy broth (TSB, Sigma-Aldrich 22092) at 37°C with 100 rpm agitation supplemented with appropriate antibiotics. Solid cultures were grown on tryptic soy agar plates (TSA) with appropriate antibiotics at 37°C.

### Cloning

All manipulations with DNA were performed following standard techniques. All cloned inserts and plasmids were sequenced to confirm integrity. For further details on plasmids and primers see Tables 2 and 3.

**Table 2:** Plasmids used in this study.

| Plasmid | Description | Reference |
| --- | --- | --- |
| pNPTS138 | Cloning suicide vector encoding *sacB*, Kan<sup>R</sup> | Kind gift of U. Jenal (Biozentrum of the University of Basel, CH)) |
| pNPTS138_Δ*bspD* (MAR135) | Suicide vector for deletion of *bspD* from *B. abortus* | This study |
| pTNS2 | Helper plasmid for miniTn7 genomic insertion | (40) |
| pUC18T-mini-Tn7T | To clone pJS203 (*tn7*::*P<sub>tet</sub>*:empty) | (40) |
| pJC44 | Template for amplification of *kanR*, *P<sub>aphT</sub>*, *dsRed* for pJS203 | (8) |
| pJS140 | pBBR-derivative with pTetAR2, *dsRed* under *P<sub>aphT</sub>*, Cm<sup>R</sup> | This study |
| pJS203 (*tn7*::*P<sub>tet</sub>*:empty) | Derivative of pUC18T-mini-Tn7T, *dsRed*, Kan<sup>R</sup>, *P<sub>TetAR</sub>* | This study |
| pMK27 (*tn7*::*eipAbspD*) | Derivative of pJS203, expression of *eipA* and *bspD* from native promoters | This study |
| pMK35 (*tn7*::*eipA*) | Derivative of pMK27, expression of *eipA* from native promoters | This study |

**Table 3:**
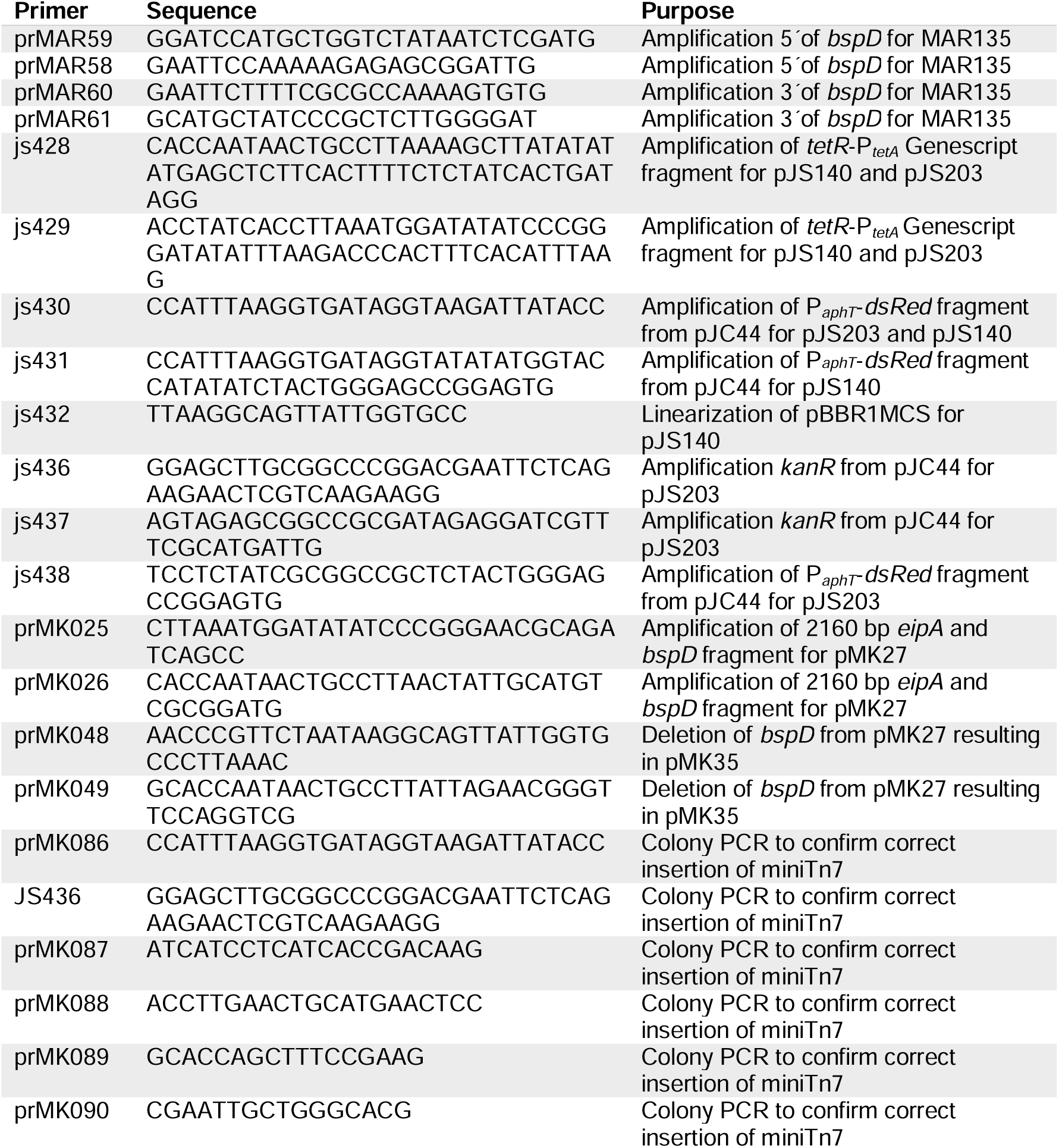
Primers used in this study.

### *Brucella* in-frame deletion mutant and complementation

*bspD* was deleted in frame by double homologous recombination using the *sacB* allelic exchange suicide vector pNPTS138, which confers kanamycin resistance and sucrose sensitivity upon insertion. Excision of the insertion was counter selected on 5% sucrose TSA plates. For further details, see (39). The miniTn7 vector pJS203 is a derivative of pUC18T-mini-Tn7T and was used for complementation of the Δ*bspD* mutant. The *kanR* cassette was amplified from pJC44 using primers js436 and js437. The P*_aphT_*-*dsRed* fragment was amplified from pJC44 with primers js430 and js438. The 705 bp *tetR*-P*_tetA_*fragment was ordered as synthetic fragment from Genescript and amplified with primers js428 and js429. All 3 fragments were then ligated into pUC18T-mini-Tn7T digested with EcoRI and XmaI using the infusion kit (Takara). The deletion of *bspD* was complemented by cloning of a 2160 bp long sequence encoding *bspD* and its upstream region including *eipA* and its 5’ non-coding region harboring the CtrA-controlled promoter (22) into pJS203 after removal of the bidirectional promoter P*_tetA_* and *tetR* through digestion with SmaI and SacI. For the complementation control *bspD* and its 5’ non-coding region were deleted from pMK27 resulting in a plasmid containing a 996 bp fragment encoding *eipA* and its 5’ non-coding region. The plasmids were finally conjugated into *Brucella* using tri-parental mating with pTNS2 as helper plasmid to ensure proper integration of the pUC18T-mini-Tn7T derivatives as described in (40). Proper integration into the genome at a secondary locus at the *att*Tn7 site between *glmS* and *recG* was confirmed by colony PCR and sequencing of colony PCR products with the primers listed in Table 3. Detailed description of plasmids and primers can be found in Tables 2 and 3, respectively.

### Growth curve, CFU plating, and analysis of bacterial morphology

Pre-cultures were grown over night to mid-exponential phase (OD_600nm_ of 1–1.4) in 3 ml TSB. 150 μl TSB with or without indicated supplements per strain in a 96 well plate (Falcon, 353072) were inoculated at OD_600nm_ of 0.08 (1.334x10^5^ CFU ml^-1^). The plate was sealed with a gas permeable seal (Azenta Life Sciences 4ti-0516/96*) and incubated at 37°C and 237 cpm for indicated times in a Synergy H1 plate reader (Agilent BioTek). OD_600nm_ was measured every 10 min. When needed, CFU ml^-1^ were enumerated at the endpoint. 70 µl of culture were centrifuged at 12000 rpm for 5 min to pellet bacteria. Bacteria were then resuspended in 140 μl PBS. 70 μl of the resuspension were used for dilution plating. For this, the samples were serially diluted 1:5 in a 96-well plate with PBS as diluent and 9 μl of each dilution were plated on TSA plates. Plates were incubated at 37°C for at least 3 days before counting and enumeration of CFU ml^-1^. To investigate bacterial morphology bacteria were harvested as described above. 500 μl 4% para-formaldehyde (PFA) were added to each sample to fix bacteria. After 20 min of incubation at room temperature PFA was removed by centrifugation and samples were washed once with PBS and resuspended in appropriate PBS volumes. Samples were spotted on 1.5% agarose-PBS pads, covered with a cover glass, and imaged using a Nikon Eclipse Ti2 with a Hamamatsu ORCA-Flash4.0 V3 Digital CMOS camera (C13440-20CU) using a 100x objective (Nikon Plan Apo Lambda 100x Oil Ph3 DM (MRD31905)). Images were analyzed using CellProfiler (version 2.2.0) (41) as described below.

### Analysis of bacterial morphology using CellProfiler

DsRed-expressing bacteria were identified as primary objects with a typical diameter of 8 to 50 pixel units using the “Global threshold strategy” with the Otsu method and “Two classes thresholding”. The threshold smoothing scale was set to 1.3488 with 1.0 as threshold correction factor and 0.0 and 1.0 as lower and upper bounds on the threshold. Clumped objects were distinguished by their intensity and dividing lines were drawn accordingly. The size of the smoothing filter for declumping as well as the minimum allowed distance between local maxima were analyzed automatically. The area of each object (“areashape_area”) was measured as a proxy for bacterial size based on the identified primary objects. Output files were one comma-separated values (csv) file per object category containing the selected measurements.

### Stress assay on plate

Stress plate assays were performed as described previously (22) with some adjustments. Bacteria were grown to stationary phase (OD_600nm_ of 3–3.6) in TSB. They were harvested by centrifugation and resuspended in sterile PBS to an OD_600nm_ of 0.1 (1.67x10^8^ CFU ml^-1^). The resuspensions were then serially diluted as described above. Finally, 9 µl of each dilution were plated on plain TSA plates and TSA plates containing either additional 200 mM NaCl (286 mM total NaCl), 2 mM EDTA, or 5 µg/ml ampicillin. Plates were incubated at 37°C for at least 3 days before counting and calculating CFU ml^-1^.

### Mammalian cell culture

The murine macrophage cell line RAW264.7 (ATCC TIB-71TM) was grown at 37°C with 5% CO_2_ in Dulbeccós Modified Eagle Medium + GlutaMAX (DMEM, Gibco 61965–026) supplemented with 10% heat-inactivated fetal calf serum (FCS, Gibco 10270).

### Infection of RAW264.7 macrophages and imaging

One day before infection 1x10^5^ cells per ml were seeded in 96-well plates (Corning 3904). Bacteria were grown to an OD_600nm_ of 1–1.4 (mid-exponential phase) in TSB. Bacteria were then diluted in DMEM/10% FCS to a multiplicity of infection (MOI) of 50 and added to the macrophages. Plates were centrifuged at 400 x g for 10 min at room temperature prior to incubation at 37°C with 5% CO_2_. 2 h post infection (hpi) the infected cells were washed twice and further incubated in DMEM/10% FCS containing 100 μg/ml gentamicin (GEN, Sigma-Aldrich G1397) to remove extracellular bacteria. Samples were fixed 27 hpi. Before fixation, samples were washed 3 times with PBS to remove detached cells and free bacteria. Samples were fixed with 3.7% para-formaldehyde (PFA, Sigma, F1635) in 0.2 M HEPES (pH 7.2– 7.4, Gibco 15630080) for 20 min at room temperature. Following incubation, samples were washed three times with PBS and cell nuclei were stained with DAPI (1 μg/ml in PBS, Sigma-Aldrich D9542) for 15 min at room temperature. Stained samples were washed three times with PBS prior to imaging. Microscopy was performed with Molecular Devices ImageXpress automated microscopes as described in (32). Nuclei were imaged using DAPI, Bacteria were identified by dsRed.

### CellProfiler analysis of infected RAW264.7 cells

Images were analyzed using CellProfiler (version 2.2.0) (41) as described in (32) with some adjustments. In brief, illumination was corrected using an illumination function based on the background method to adjust uneven illumination. Images were then smoothed using a Gaussian method with 100-filter size. To remove the signal originating from *Brucella* DNA in the DAPI channel the dsRed image was subtracted from the DAPI image. Objects were segmented and measured as follows: DAPI-stained nuclei were detected as primary objects using the Automatic threshold strategy. Clumped objects were distinguished using their Intensity and dividing lines were drawn according to their shape using a smoothing filter of 5. Nuclei were filtered based on their size features using a minimum measurement value of 0.1 for AreaShape with the Measurement FormFactor. DsRed-expressing bacteria were detected as microcolonies with a typical diameter of 6 to 30 pixels as primary objects using the “Adaptive threshold strategy” followed by a “Background method”. Smoothing was done “Automatic” with a “Threshold correction factor” of 1.35 with 0.04 lower and 1.0 upper bounds on threshold. Clumped objects were distinguished and divided using their “Intensity” with a smoothing filter of 8. The cell body was estimated by expansion of the nuclei by 9 pixels. Bacteria partially or entirely within the estimated cell body were scored as 1. The number of cells with microcolonies and the size of the microcolonies (“areashape_area”) were reported for each image. To set boundaries for the identification of microcolonies the typical diameter of *B. abortus* wild-type vs. Δ*virB9* mutant were determined from the measured area using dd = 2(√A/√π) (dd – typical diameter, A – area) (Fig. S5). Data from the same condition from separate wells were pooled. Output files were one comma-separated values (csv) file per object category containing the selected measurements.

### Identification of BspD orthologs, phylogenetic analysis, and synteny

The synteny for chosen species was built using the SyntTax browser (https://archaea.i2bc.paris-saclay.fr/SyntTax/) (31).

The species tree of life based on the species phylogeny of the hierarchical orthologous groups (HOGs) was recovered from the OMA browser (https://omabrowser.org/oma/current/) (42) and simplified to representative species of the genus Rhizobiales. The presence or absence of *bspD* (BAB1_1611) and *eipA* (BAB1_1612) orthologs was based on their respective HOGs retrieved from OMA (42). Co-conservation with *ctrA* and *chpT* in the same locus was deduced from SyntTax (31). The presence or absence of annotated VirB-T4ASSs was inferred from the SecReT4 database (https://bioinfo-mml.sjtu.edu.cn/SecReT4/index.php) (43). The different lifestyles were inferred from literature.

Sequences of BspD homologues were retrieved from the OMA browser (42) based on the OMA Group MVEPIMY of BAB1_1611 from *B. abortus* 2308. 25 orthologues from different members of the Rhizobiales were taken for the phylogenetic analysis presented in Fig. S1 and S2. There were no orthologues detected outside of the Rhizobiales. Species and strain names are listed in Table S1.

Protein sequences were aligned with Geneious Prime 2019.0.4 using Geneious Alignment with standard settings (Fig. S1 and supplementary data file 1). The maximum likelihood phylogeny presented in Fig. S2 was built with Geneious using PhyML with standard settings and bootstrap 100 (44).

### Statistical analysis

Graphs were created with GraphPad Prism 9. Statistical analysis was performed using GraphPad Prism with one-way ANOVA with Tukeýs multiple comparison test with *p≤0.0371, **p≤0.0066, ***p≤0.0003, and ****p<0.0001. When indicated outlier were removed using the ROUT method with Q=10%. The number of independent replicates is indicated in the figure legends as n.

## Acknowledgments

This work was supported by the Swiss National Science Foundation (SNSF, www.snf.ch) grant 310030B_201273 (to C.D.). We thank Mariana Martins for contributing the Δ*bspD* mutant.

