## Supplemental Figures and Tables for "The putative Type 4 secretion system effector BspD is involved in maintaining envelope integrity of the pathogen *Brucella*"

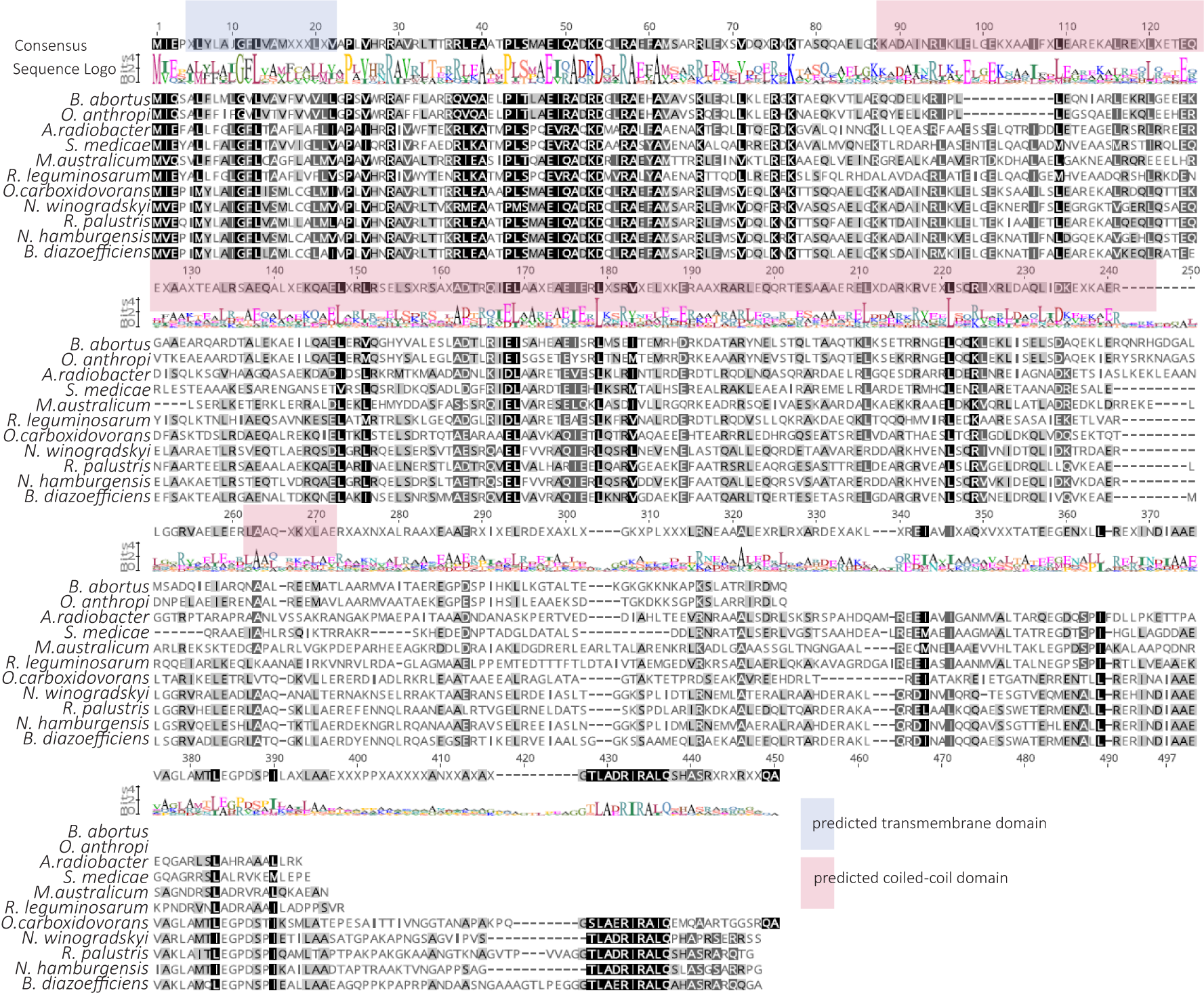

**Fig. S1: Protein sequence alignment of BspD orthologues of diverse species within the Rhizobiales.** Multi-sequence alignment of protein sequences of BspD from the following species (accession numbers indicated in brackets): *B. abortus* (BAB1_1611), *O. anthropi* (A0A6I0CU19), *A. radiobacter* (B9J8Q3), *S. medicae* (A0A508WYJ9), *M. australicum* (L0KKH9), *R. leguminosarum* (Q1MCS8), *O. carboxidovorans* (WP_012564253), *N. winogradskyi* (Q3SVA9), *R. palustris* (Q6N9C7), *N. hamburgensis* (Q1QI36), and *B. diazoefficiens* (A0A2A6MP69). Predicted transmembrane and coiled-coil domain from BspD of *B. abortus* highlighted in blue and pink, respectively. For a full aligment see supplementary data file 1.

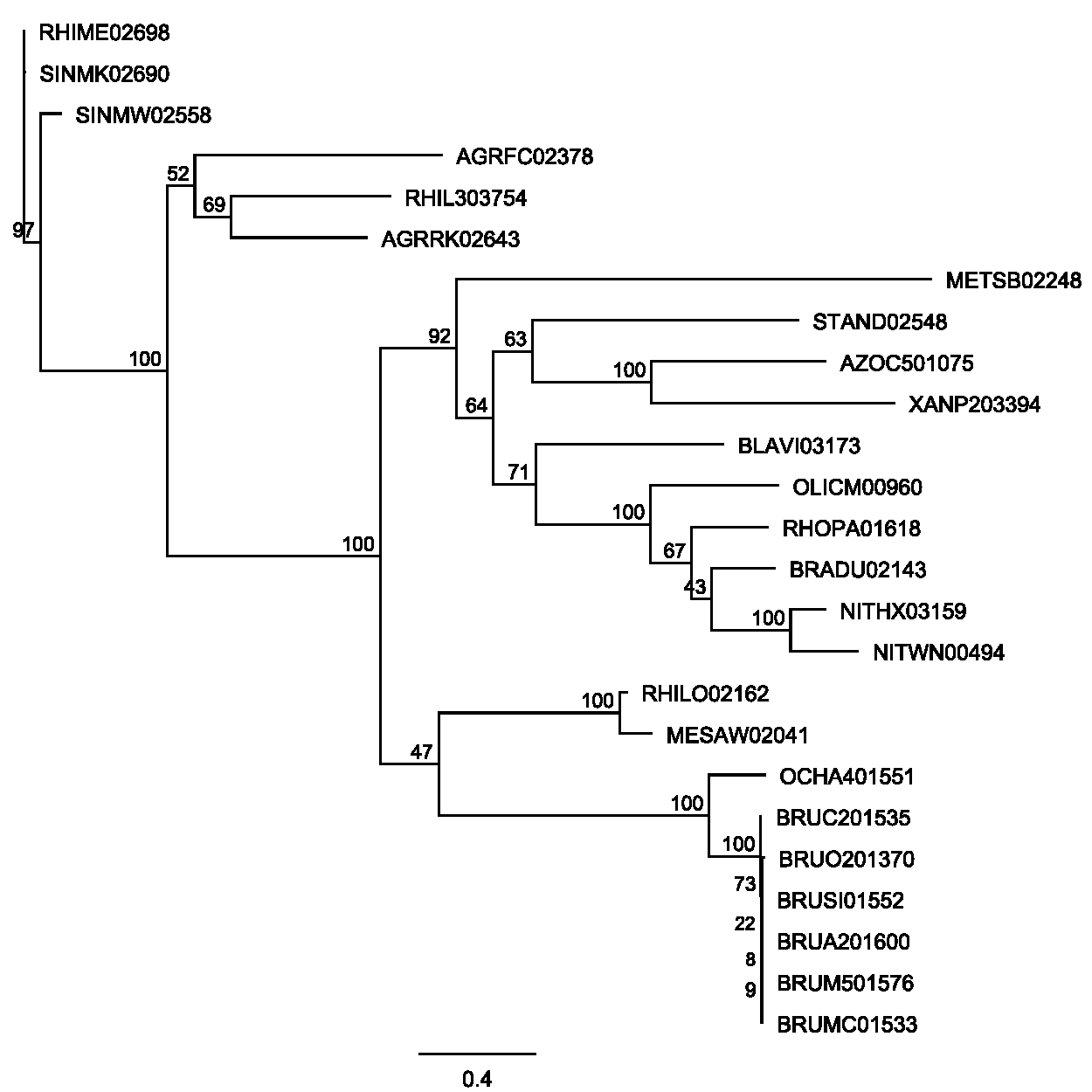

**Fig. S2: Phylogenetic tree of BspD.** Geneious PhyML tree based on alignment of 25 sequences from OMA group MVEPIMY. Bootstrap values above 50 are shown. Line at bottom depicts 0.4 substitutions per site. Abbreviations are as follows: BRUO201370 - *Brucella ovis* A0A0H3ASC5, BLAVI03173 - *Blastochloris viridis* A0A0H5BNY9, BRADU02143 - *Bradyrhizobium diazoefficiens* A0A2A6MP69, OCHA401551 - *Ochrobactrum anthropic* A0A6I0CU19, A0A7Y0XP19 - *Agrobacterium fabrum* A0A7Y0XP19, RHIME02698 - *Rhizobium meliloti* A0A222HNP4, SINMW02558 - *Sinorhizobium medicae* A0A222HNP4, XANP203394 - *Xanthobacter autotrophicus* A7IL20, AZOC501075 - *Azorhizobium caulinodans* A8HQZ7, BRUC201535 - *Brucella canis* A9M704, BRUSI01552 - *Brucella suis* B0CI72, METSB02248 - *Methylocella silvestris* B8EIB6, AGRRK02643 - *Agrobacterium radiobacter* B9J8Q3, BRUM501576 - *Brucella melitensis* BM590, BRUMC01533 - *Brucella microti* C7LDI5, STAND02548 - *Starkeya novella* D7A4M7, MESAW02041 - *Mesorhizobium australicum* L0KKH9, RHIL303754 - *Rhizobium leguminosarum bv. Viciae* Q1MCS8, NITHX03159 - *Nitrobacter hamburgensis* Q1QI36, BRUA201600 - *Brucella abortus* Q2YQA7, NITWN00494 - *Nitrobacter winogradskyi* Q3SVA9, RHOPA01618 - *Rhodopseudomonas palustris* Q6N9C7, RHILO02162 - *Mesorhizobium japonicum* Q98HM0, SINMK02690 - *Sinorhizobium meliloti* Sinme_2777, OLICM00960 - *Oligotropha carboxidovorans* WP_012564253

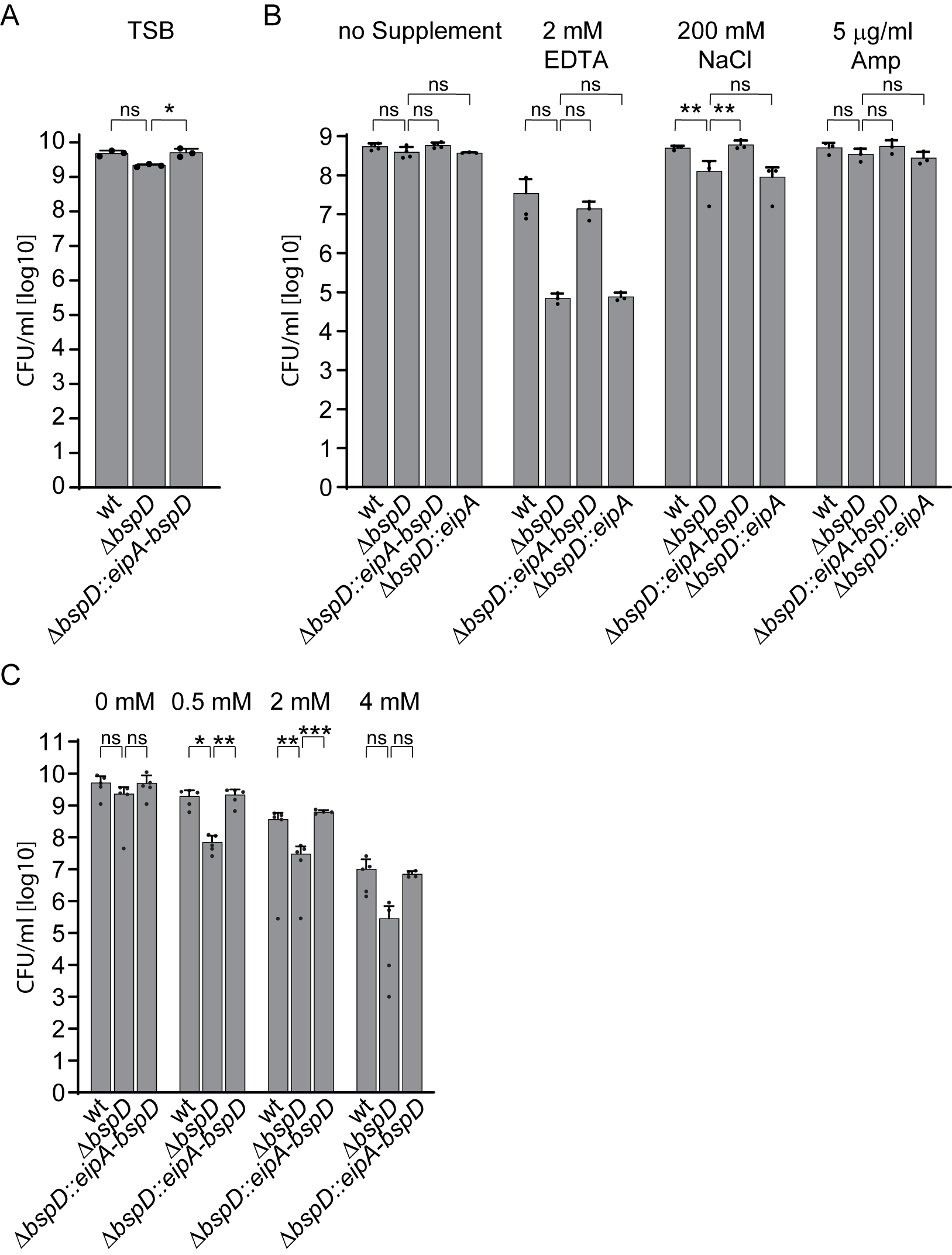

**Fig. S3: CFU enumerations of stress plate assay and growth curves.** **(A)** CFU enumeration after growth in TSB for 96 h. Corresponding to Fig. 2C. **(B)** CFU enumeration after growth on TSA supplemented with indicated stressors for at least 2-3 days at 37 °C. Corresponding to Fig. 3A. **(C)** CFU enumeration at endpoint after growth in TSB supplemented with indicated concentrations of EDTA for 96 h. Corresponding to Fig 3C. Dots represent individual experiments +SD. Statistical analysis was performed with ordinary one-way ANOVA followed by Tukey’s multiple comparison test to Δ*bspD*.

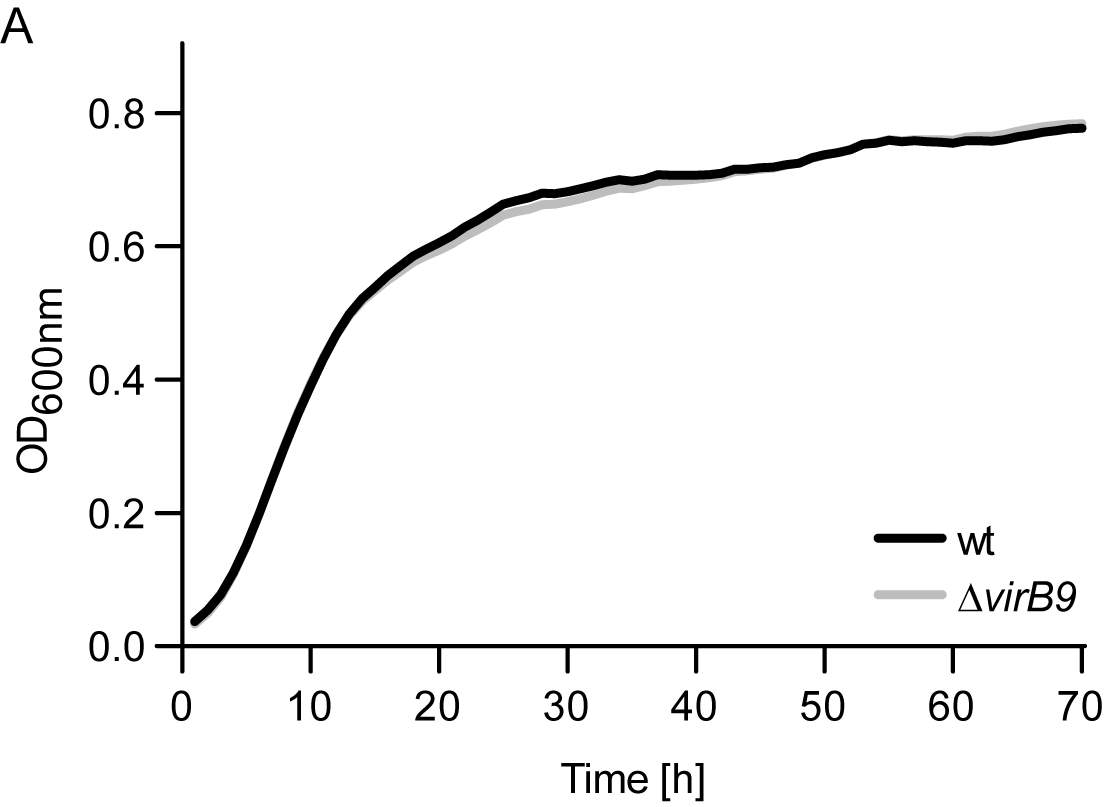

**Fig. S4: Wild-type and ∆*virB9* display a similar growth kinetics in TSB.** **(A)** Optical density of indicated *Brucella* strains grown in 150 µl TSB in a 96 well plate over 96 h with a starting OD_600nm_ of 0.08. The OD_600nm_ was measured every 10 min in a Synergy H1 plate reader. Lines depict means of 2 independent experiments with a 2nd order smoothing of 15 neighbors applied in GraphPad Prism 9.3.1. n=2 with 3 technical replicates each.

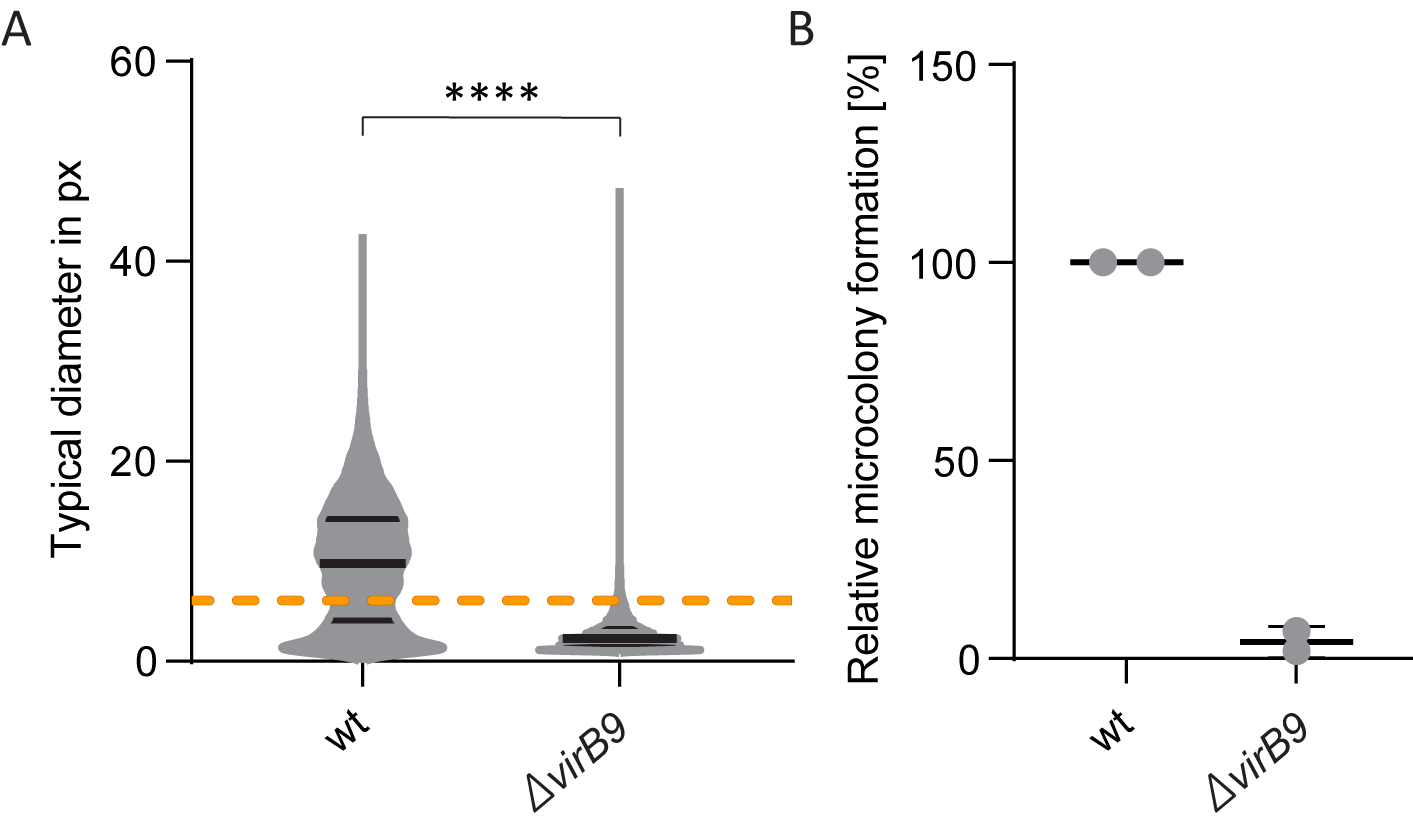

**Fig. S5: Definition of thresholds for the identification of microcolonies in infected RA264.7 macrophages.** **(A)** RAW264.7 macrophage-like cells were infected with an MOI of 50 for 27 hpi. After fixation and staining of host cell nuclei with DAPI, samples were imaged using a Molecular Devices ImageXpress microscope. Images were analyzed using cell profiler. The typical diameter of a circle was calculated from measured areas in pixel units (px) with the formula dd = 2(√A/√π). Orange line – 95% percentile of the ∆*virB9* population at 6.077. Black – mean. n=1 with over 17000 events analyzed. **(B)** RAW264.7 cells were infected and processed as in (A). The relative microcolony formation was measured using a CellProfiler pipeline with lower threshold for identification of microcolonies set at 6 pixels and upper threshold at 30 pixels based on the experiment presented in (A). n=2.

**Table S1: Strains used for the phylogenetic analysis of BspD orthologues of OMA group MVEPIMY**

| Accession Number | Protein ID | Species Name | Strain Identifier |
| --- | --- | --- | --- |
| A0A0H3ASC5 | BRUO201370 | *Brucella ovis* | ATCC 25840 |
| A0A0H5BNY9 | BLAVI03173 | *Blastochloris viridis* | - |
| A0A2A6MP69 | BRADU02143 | *Bradyrhizobium diazoefficiens* | JCM 10833 |
| A0A6I0CU19 | OCHA401551 | *Ochrobactrum anthropi* | ATCC 49188 |
| A0A7Y0XP19 | AGRFC02378 | *Agrobacterium fabrum* | ATCC 33970 |
| A0A222HNP4 | RHIME02698 | *Rhizobium meliloti* | 1021 |
| A0A508WYJ9 | SINMW02558 | *Sinorhizobium medicae* | WSM419 |
| A7IL20 | XANP203394 | *Xanthobacter autotrophicus* | ATCC BAA-1158 |
| A8HQZ7 | AZOC501075 | *Azorhizobium caulinodans* | ATCC 43989 |
| A9M704 | BRUC201535 | *Brucella canis* | ATCC 23365 |
| B0CI72 | BRUSI01552 | *Brucella suis* | ATCC 23445 |
| B8EIB6 | METSB02248 | *Methylocella silvestris* | DSM 15510 |
| B9J8Q3 | AGRRK02643 | *Agrobacterium radiobacter* | ATCC BAA-868 |
| BM590 | BRUM501576 | *Brucella melitensis* | M5-90 |
| C7LDI5 | BRUMC01533 | *Brucella microti* | CCM 4915 |
| D7A4M7 | STAND02548 | *Starkeya novella* | ATCC 8093 |
| L0KKH9 | MESAW02041 | *Mesorhizobium australicum* | HAMBI 3006 |
| Q1MCS8 | RHIL303754 | *Rhizobium leguminosarum bv. viciae* | 3841 |
| Q1QI36 | NITHX03159 | *Nitrobacter hamburgensis* | DSM 10229 |
| Q2YQA7 | BRUA201600 | *Brucella abortus* | 2308 |
| Q3SVA9 | NITWN00494 | *Nitrobacter winogradskyi* | ATCC 25391 |
| Q6N9C7 | RHOPA01618 | *Rhodopseudomonas palustris* | ATCC BAA-98 |
| Q98HM0 | RHILO02162 | *Mesorhizobium japonicum* | LMG 29417 |
| Sinme_2777 | SINMK02690 | *Sinorhizobium meliloti* | AK83 |
| WP_012564253 | OLICM00960 | *Oligotropha carboxidovorans* | OM4 |
